# Proteomic and transcriptomic analysis of *Microviridae* φXI74 infection reveals broad up-regulation of host membrane damage and heat shock responses

**DOI:** 10.1101/2020.10.26.355149

**Authors:** Bradley W Wright, Dominic Y Logel, Mehdi Mirzai, Dana Pascovici, Mark P Molloy, Paul R Jaschke

## Abstract

Measuring host-bacteriophage dynamics is an important approach to understanding bacterial survival functions and responses to infection. The model *Microviridae* bacteriophage φX174 is endemic to the human gut and has been studied for over seventy years but the host response to infection has never been investigated in detail. To address this gap in our understanding of this important interaction within our microbiome we have measured host *Escherichia coli* C proteomic and transcriptomic response to φX174 infection. We used mass spectrometry and RNA-seq to identify and quantify all 11 φX174 proteins and over 1,700 *E. coli* proteins, enabling us to comprehensively map host pathways involved in φX174 infection. Most notably, we see significant host responses centered on membrane damage and remodeling, cellular chaperone and translocon activity, and lipoprotein processing, which we speculate is due to the peptidoglycan-disruptive effects of the φX174 lysis protein E on MraY activity. We also observe the massive upregulation of small heat-shock proteins IbpA/B, along with other heat shock pathway chaperones, and speculate on how the specific characteristics of holdase protein activity may be beneficial for viral infections. Together, this study enables us to begin to understand the proteomic and transcriptomic host responses of *E. coli* to *Microviridae* infections and contributes insights to the activities of this important model phage.

**IMPORTANCE:** A major part of the healthy human gut microbiome are the *Microviridae* bacteriophage, exemplified by the model φX174 phage. Although much has been learned from studying φX174 over the last half century, until this work, the *E. coli* host response to infection has never been investigated in detail. We reveal the proteomic and transcriptomic pathways differentially regulated during the φX174 infection cycle, and uncover the details of a coordinated cellular response to membrane damage that results in increased lipoprotein processing and membrane trafficking, likely due to the phage antibiotic-like lysis protein. We also reveal that small heat shock proteins IbpA/B are massively upregulated during infection and that these holdase chaperones are highly conserved across the domains of life, indicating that reliance on them is likely widespread across viruses.

## INTRODUCTION

Successful phage reproduction relies on suppression and evasion of host defense responses (1), and to varying extents, requisitioning host machinery (2). Temporal infection studies employing the use of transcriptomics (3, 4) and proteomics (5–11) have revealed insights into the establishment of infection and utilization of host machinery. In particular, the use of modern mass spectrometry work-flows and instruments to identify and quantify thousands of proteins, is an invaluable tool for the study of complex biological systems especially when paired with complimentary RNA-sequencing (RNA-seq) methods. Currently, most phage-host interactions, even of model systems, have not yet been characterized using these methods.

Bacteriophage φX174 is a member of the family *Microviridae*, which make up a group of small icosahedral viruses encoded by a single-stranded DNA genome. The small 5,386 nucleotide genome of φX174 containing 11-protein encoding genes makes it a tractable target for proteomic and transcriptomic studies (12, 13). *Microviridae* have been used as model systems for DNA sequencing, genome engineering (14, 15), evolution (16), icosahedral virus packaging (17, 18), and novel virus infection mechanisms (19, 20). Furthermore, φX174 and other *Microviridae* are universal members of the human gut microbiome (21, 22), although their precise role is not well understood.

In this investigation we applied modern proteomics and complementary transcriptomics to map for the first time the temporal response of *Escherichia coli* NCTC122 (C122) to phage φX174 infection. Our detailed measurements of protein and RNA revealed the heat shock chaperone network is selectively up-regulated during infection, along with many proteins involved in membrane repair and remodeling, and membrane transporters. Most notably, we observed massive up-regulation of small heat-shock proteins (sHSPs) IbpA and IbpB at both the protein and RNA level. This work lays the foundation for further studies of *Microviridae* infections and their interplay within a range of more natural host strains.

## RESULTS

### φX174 infection results in moderate changes to the host proteome and transcriptome just prior to lysis

To measure the host response to φX174 infection, we first used a mass spectrometry proteomic work-flow to compare φX174 infected *E. coli* C122 and mock-infected cells at five time-points (0, 15, 30, 60, and 75-minutes post-infection). We found no differences between the host proteome expression and mock-infections until 60 and 75-minutes, corresponding to the point at which we observed lysis under our conditions (Figure S1A). Mass spectrometry analysis identified 2,184 host proteins (Supplementary File S1), representing ~57 % of the *E. coli* C122 proteome database. Of the identified proteins, we confidently measured the expression of 1,752 host proteins and all 11 φX174 proteins. To further validate our proteomics work we also measured the transcriptional response of the host during late lysis period and mapped reads to 85% of *E. coli* protein-encoding genes and 100% of φX174 genes. Furthermore, during infection, phage-specific reads accounted for 18 % of the total mapped reads.

During late-infection we measured at the protein-level 255 up-regulated and 54 down-regulated proteins (Figure 1A and Supplemental File S1), while at the RNA-level 203 genes were found to be up-regulated and 325 genes were down-regulated (Figure 1B and Supplementary Figure S2). The *ibpA* and *ibpB* small heat shock proteins, which had fold-changes (FCs) of +8.7 / +27.1, and +15.6 / +38.6 at the protein/RNA-levels respectively, were similar in magnitude and significance to the φX174 proteins (Figure 1C). Notably, we did not observe any significant changes to small RNAs within *E. coli* C122 due to φX174 infection (Supplemental File S2).

**Figure 1:**
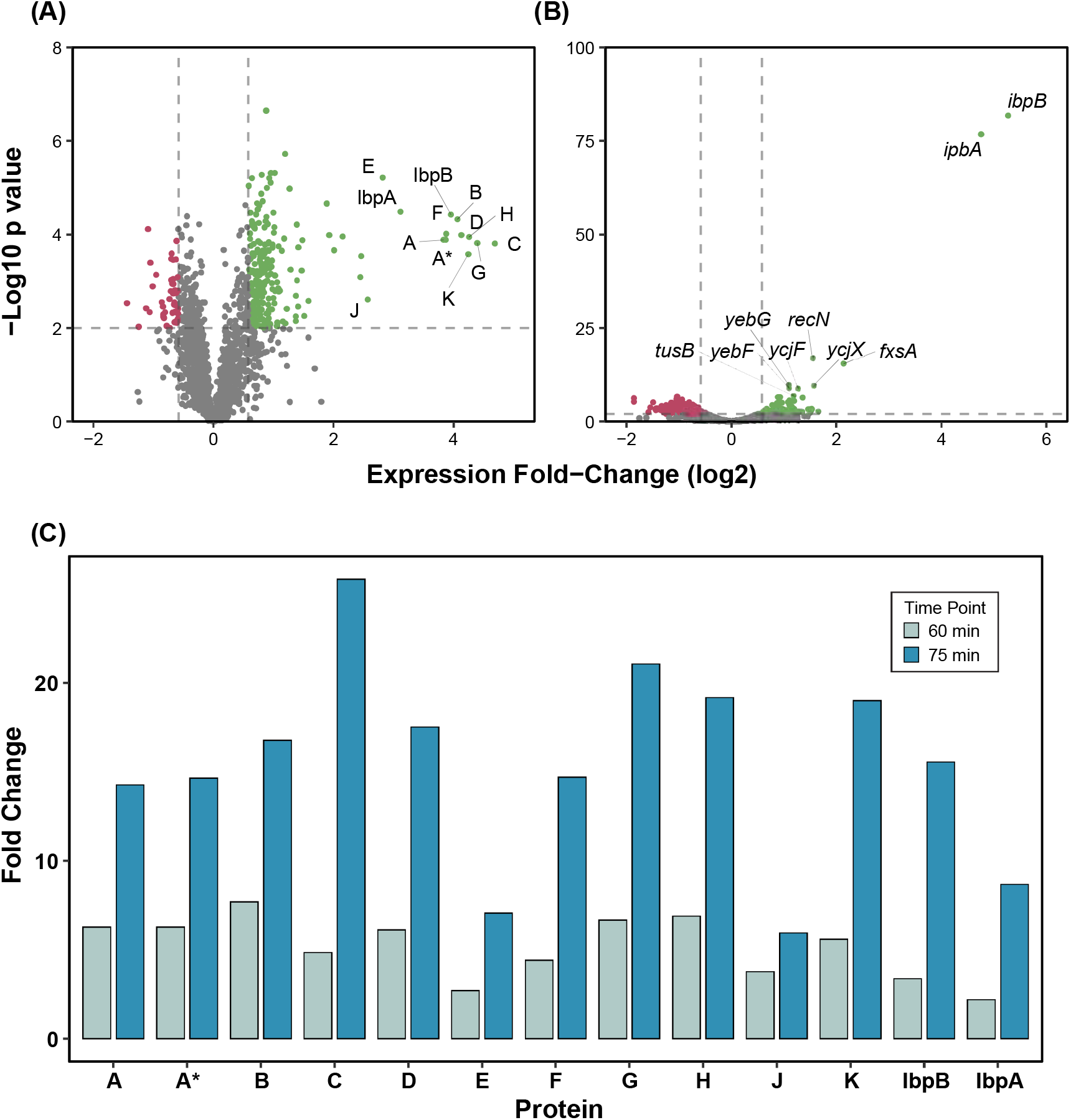
A moderate number of *E. coli* C122 genes are differentially regulated during late φX174 infection, with *ibpAB* up-regulation equal to φX174 phage genes. **(A)** Quantified proteins. FC ± 1.5 and p-value <0.025 significance threshold. **(B)** Quantified RNA. FC ± 1.5 and p-value cut-off <0.05 significance threshold. **(C)** Foldchanges for phage proteins and host proteins IbpA and IbpB at 60-minute and 75-minutes post-infection.

### Host engages in significant membrane protein up-regulation and metabolic protein down-regulation during infection

To define the broader biological impacts of φX174 infection, we assigned all 1,752 quantified host proteins, 11 φX174 proteins, and 528 differentially expressed RNA transcripts to their respective clusters of orthologous groupings (COG) using EggNOG-mapper (23). The results of this analysis showed no sample preparation enrichment bias as the “Annotated C122” and “Quantified Proteins” annotated biological function category COG distributions were comparable (Figure 2).

**Figure 2:**
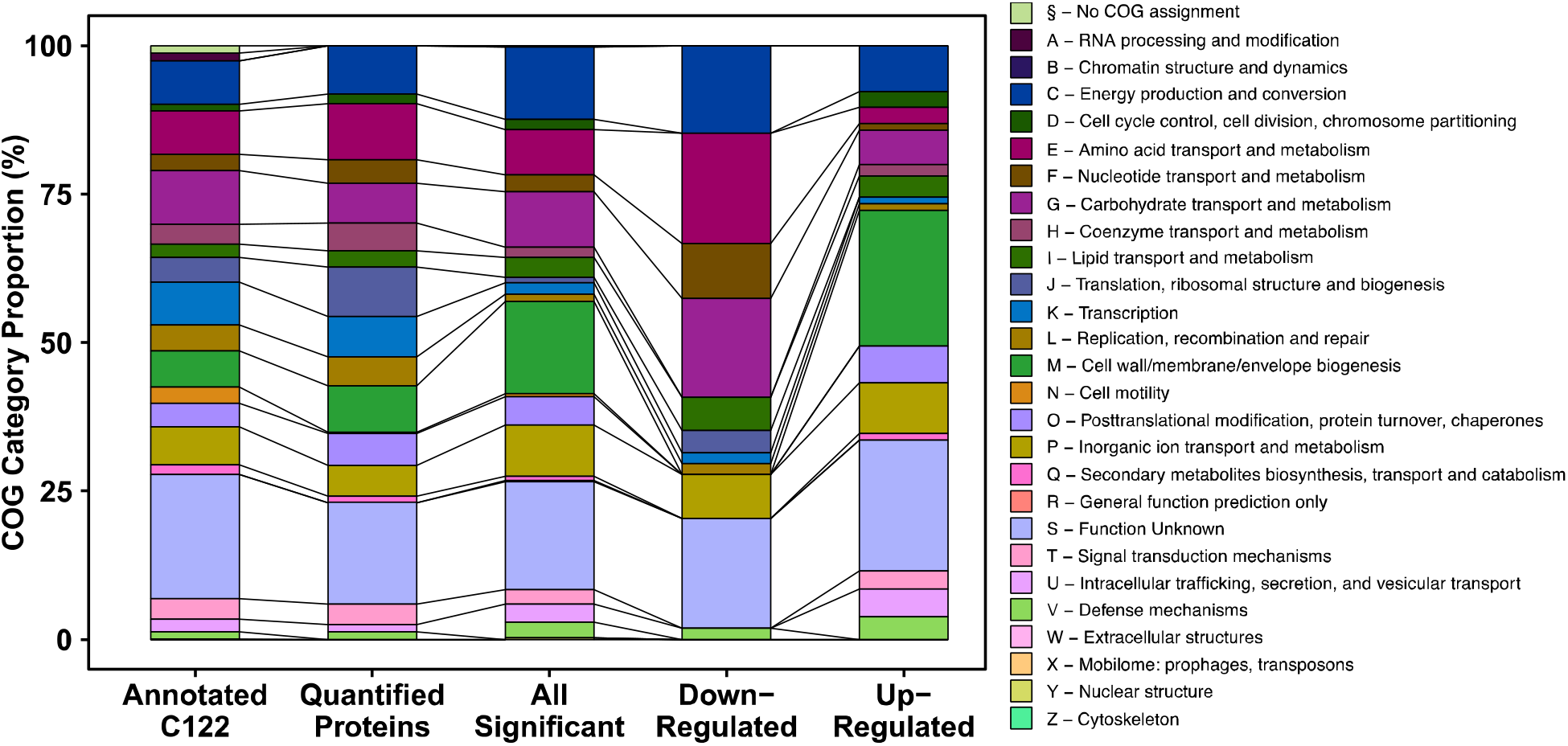
Clusters of orthologous groups (COG) distributions of measured *E. coli* C122 proteins. Annotated C122: 3,666 annotated proteins; Quantified Proteins: 1,763 proteins quantified across all conditions; All Significant: Non-redundant list of 399 significantly differentially regulated proteins across all time-points and samples; Down-regulated: 54 significantly down-regulated proteins from 75-minutes post-infection; Up-regulated: 255 significantly up-regulated proteins from 75-minutes post-infection. COGs were assigned to C122 genes using the EggNOG-mapper (23).

Within the group of 309 significantly differentially regulated proteins at 75-minutes post-infection (“All Significant”), we found that all proteins annotated with COG categories M (cell wall/membrane/envelope biogenesis) and O (Post-translational modification, protein turnover, chaperones) were up-regulated during infection (Figure 2). These category enrichments indicate there may be a host response mediating significant membrane restructuring or repair, or poor translocation of membrane bound proteins leading to their accumulation in the cytoplasm. Within the group of 54 down-regulated proteins we found the majority of these proteins (~59%) were in COG categories representative of energy and metabolism (C, E, F, and G) (Figure 2).

These findings were further reinforced with a PANTHER (24) over-representation test, whereby 167 of the 255 up-regulated proteins were mapped according to their GO cellular component annotations. We found the majority (60%) were mapped to the membrane component (p-value = 1.54E-57), and interestingly, of the 54 proteins significantly down-regulated as a result of phage infection, a significant proportion (62%) were mapped to the cytoplasm (p-value = 2.10E-04).

Corresponding with the proteomic findings, the transcriptomics showed COG category enrichment for inorganic ion transport and metabolism (P), amino acid transport and metabolism (E), and energy production and conversion (C) (Supplementary Figure S3). Additionally, the transcriptomics showed enrichment for the COG categories cell motility (N), and intracellular trafficking, secretion, and vesicular transport (U). PANTHER over-representation testing for GO cellular component terms showed significant enrichment within up-regulated genes for the pilus (6.7 fold) and the integral component of the membrane (1.6 fold). The down-regulated genes were enriched for the iron-sulfur cluster assembly complex (16.2 fold), the proton-transporting ATP synthase complex, catalytic core F(1) (13.5 fold), and the plasma membrane respiratory chain complex I (11.2 fold) (Supplementary Figure S4). All of these enriched cellular components are localized within the bacterial membrane. Membrane component enrichment was reflected in the differentially expressed genes more broadly as well, as gene products destined for the membrane accounted for 42 % and 24 % of the up- and down-regulated datasets, respectively.

To understand the interaction of φX174 protein expression and host responses to infection, we performed comparative analyses using an UpSet plot, which is a tool to visualize and group shared elements within intersecting datasets (25). We used the UpSet visualization to identify the conditions and samples in which significantly differentially regulated proteins are shared. This analysis revealed that the 60-minutes post-infection samples (60C and 60P) shared 12 differentially expressed proteins with the 75-minute post-infection samples (75C and 75P): two host proteins (IbpA, and IbpB), and all φX174 proteins except C protein (Figure 3A and Supplementary File S1).

**Figure 3:**
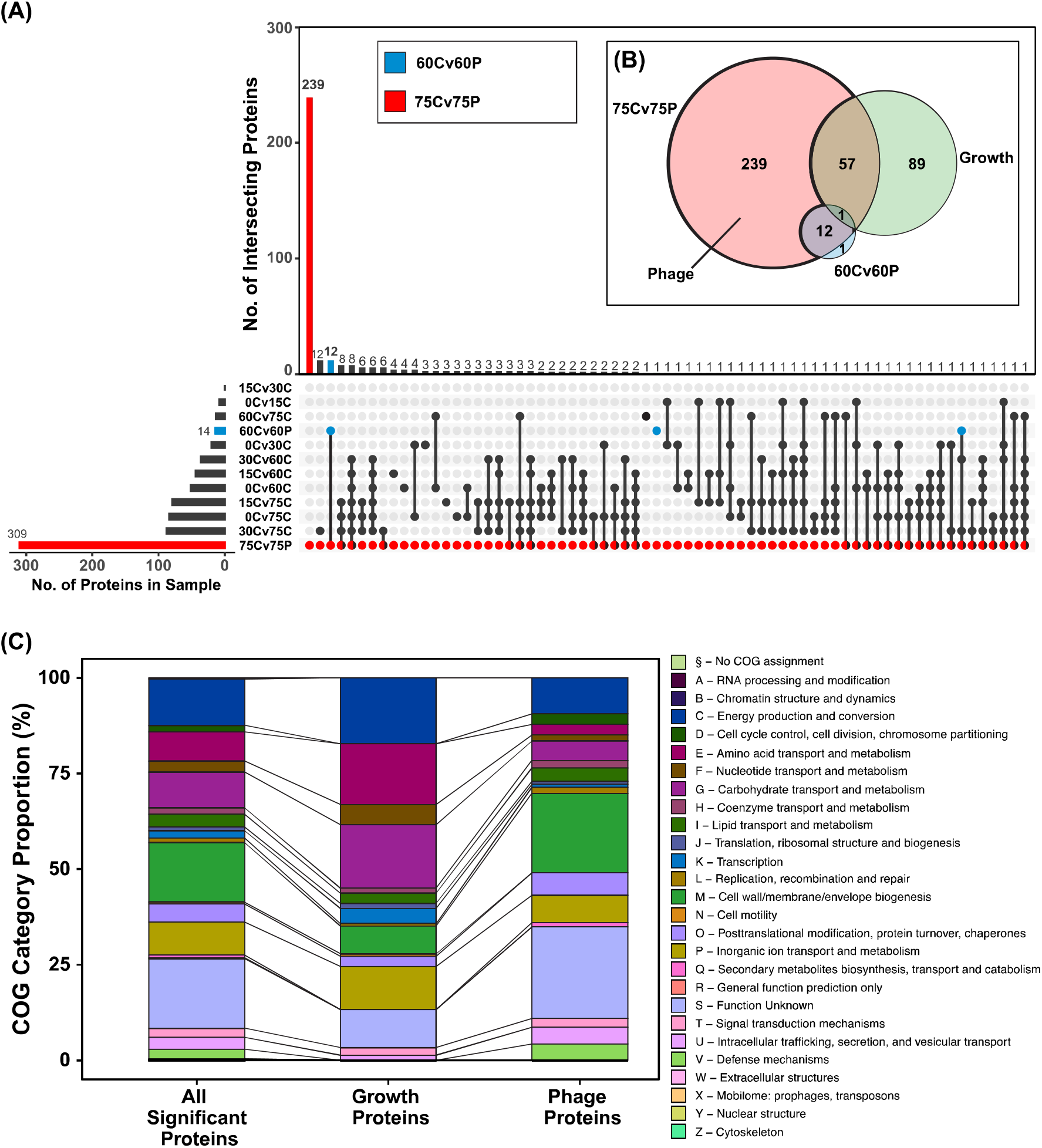
Sets of significantly differentially regulated proteins due to cellular growth and φX174 infection and their functional groupings. **(A)** UpSet visualization of sample groups analyzed by proteomics. Each set represents the group of significantly regulated proteins (FC ±1.5, p-value <0.025) extracted from comparative analyses. For example, 75Cv75P (red) is the group of significant proteins passing our stated cut-off thresholds when the 75-minute mock-infected (75C) group is compared to the 75-minute φX174-infected (75P) group. This plot allows visualization of redundant proteins found differentially regulated across multiple groups, and more importantly, facilitates the partitioning out of significant proteins relevant only to φX174 infection. Non-redundant sample size = 399 proteins. **(B)** Euler plot showing relationships between Phage, Growth, 60Cv60P, and 75Cv75P protein sets. **(C)** Clusters of orthologous groups (COG) categories of the Growth and Phage protein sets as compared to 399 significant proteins (“All Significant Proteins”). See Supplemental File S1. COGs were assigned to C122 genes using the EggNOG-mapper (23).

In contrast to the 60-minute samples, the 75-minutes post-infection samples showed large-scale changes to the host proteome, with 309 differentially regulated proteins including all 11 phage proteins and the same two host proteins, IbpA and IbpB, seen at 60-minutes (Figure 3A). Within this group of 309 proteins we wanted to identify proteins that were strictly impacted due to the phage infection and not cellular growth, which was also occurring during the phage infection. To identify proteins that changed in expression due to cellular growth alone we compared all permutations of mock-infected time-points and identified 147 proteins that were differentially regulated due to cell growth. We called this group Growth (Figure 3B). Comparing the Growth proteins to the 309 proteins differentially regulated at 75-minutes post-infection, revealed a set of 58 proteins differentially regulated due to both Growth as well as φX174 infection (Figure 3B). Removing these 58 proteins along with the 12 proteins shared with the 60-minutes post-infection sample gave a set of 239 proteins differentially regulated due solely to phage infection. We called this group Phage (Figure 3B).

Examining the biological functions enriched within the Growth protein set revealed proteins involved in metabolic activity such as energy production and conversion (C), and amino acid (E) and carbohydrate (G) transport and metabolism (Figure 3C). In contrast, Phage proteins were enriched for a distinct set of categories: cell wall/membrane/envelope biogenesis (M), intracellular trafficking, secretion, and vesicular transport (U), and defense mechanisms (V) (Figure 3C).

### Disruption to peptidoglycan synthesis and up-regulation of membrane tethering lipoproteins associated with φX174 protein E expression

Looking deeper into the COG category M (cell wall/membrane/envelope biogenesis) we found proteins from the outer membrane lipoprotein maturation and transportation pathway were significantly up-regulated. Typically, a protein is translocated across the inner membrane from the cytoplasm by the Sec pathway, or via the twin-arginine translocation (TAT) pathway. We did not observe differential regulation of the TAT pathway, but did see the up-regulation of all components of the cytoplasmic membrane complex of the Sec pathway (SecYEG, FCs +1.6, +1.8, and +1.8, respectively) and the accessory protein YidC (FC +1.8) (26) (Figure 4). PANTHER overrepresentation analysis of proteins identified as up-regulated from the Phage proteins set (Supplemental File S1) highlighted that the cell envelope Sec protein transport complex is significantly over-represented (p-value < 0.0002) during infection. This complex facilitates the translocation of nascent unfolded proteins into the periplasm in an ATP-dependent fashion (27). We did not identify differential expression of SecA, an ATPase that interacts with SecYEG and SecB (28), nor the significant differential expression of SecB itself. SecF was not observed but its binding partner, SecD, was significantly up-regulated (FC +1.7). YajC also interacts with SecDF and we saw it significantly up-regulated (FC +1.8) to a similar magnitude during φX174 infection (Figure 4).

**Figure 4:**
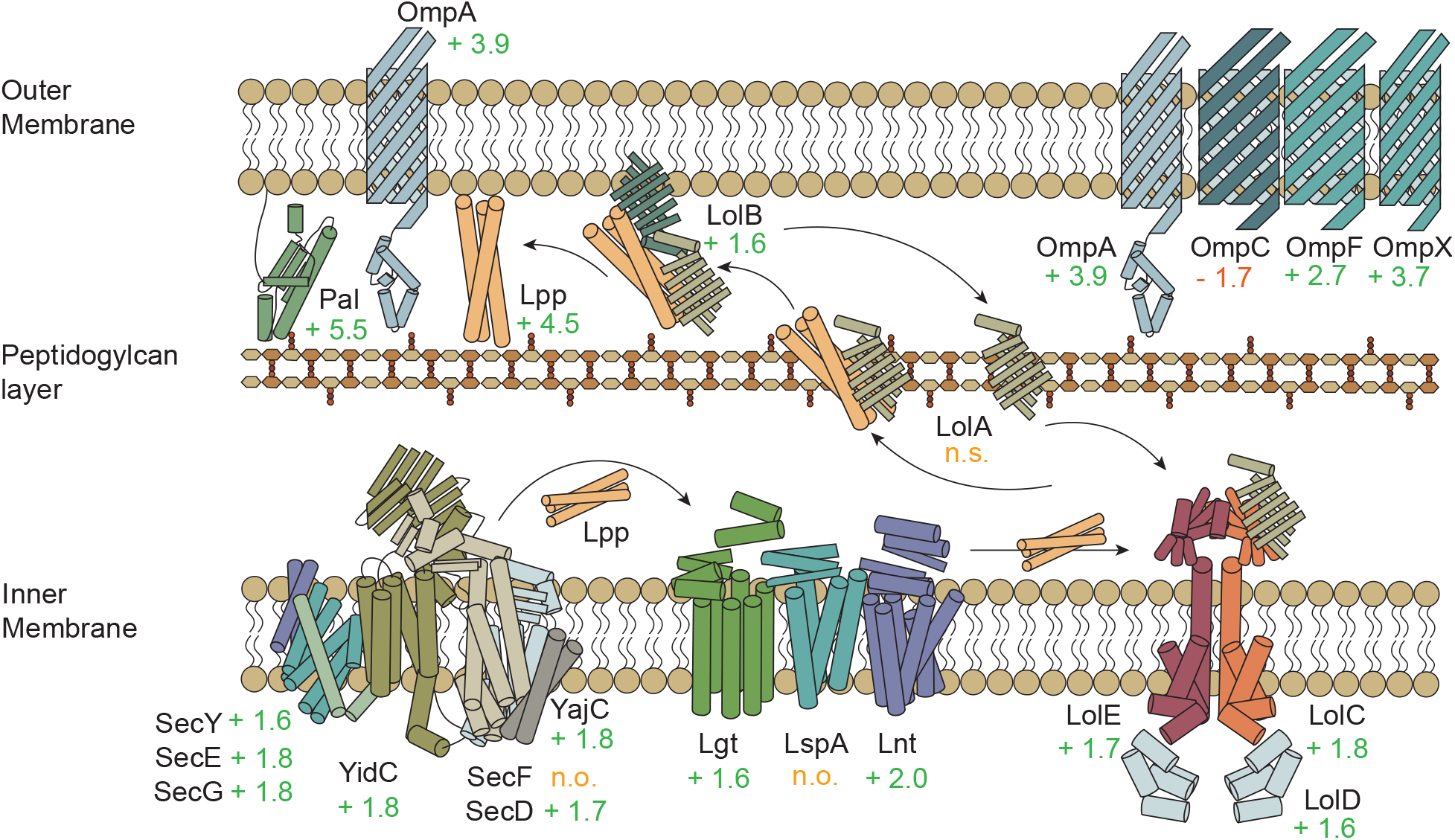
*E. coli* C122 upregulating membrane stabilizing and anchoring proteins Lpp, OmpA, and Pal, as a result of protein E membrane disruption. Up-regulation of the membrane lipoprotein maturation and transportation pathway. Depicted are nascent lipoproteins translocating across the inner membrane through the Sec pathway, followed by processing, and Lol pathway-mediated export to the outer membrane. Outer membrane proteins are depicted - including OmpA, which has its peptidoglycan anchoring role highlighted along with Lpp and Pal. Values indicate protein fold-change during infection (75P/75C protein samples), n.o: protein not observed, n.s: protein observed but not significantly altered in abundance between conditions.

After nascent lipoproteins are translocated through the inner membrane via the Sec pathway machinery, a diacylglycerol group is added to the protein by the inner membrane diacylglycerol transferase protein Lgt (29). We observed the significant upregulation of Lgt in the phage infected samples (FC +1.6) as well as the Lnt protein that subsequently performs N-terminal acylation (FC +2.0) (29), but we did not observe signal peptide cleaving protein LspA (Figure 4). The ATP-driven LolCDE inner membrane complex (FCs +1.8, +1.6, and +1.7, respectively) releases lipoproteins to the periplasmic chaperone protein LolA (expression not significantly changed) (30). From there, LolA facilitates the lipoprotein’s transfer to the outer membrane receptor LolB (FC +1.6) which mediates their integration into the outer membrane (31) (Figure 4).

The up-regulation of the lipoprotein processing and trafficking machinery was in concordance with the up-regulation of the major outer membrane lipoprotein (Lpp) in the Phage protein set (FC +4.5) (Figure 4). Lpp tethers the outer membrane to the peptidoglycan layer through a covalent linkage, facilitating structural integrity and maintenance of cell shape (32). Lpp attachment to the outer membrane is dependent on LolB (FC +1.6), which if depleted, is detrimental to the cell due to Lpp mislocalization (33). Similar to Lpp, we saw up-regulation of OmpA and Pal (FCs +3.9 and +5.5), which also function in maintenance of cell membrane integrity, and act through non-covalent interactions connecting the outer membrane and peptidoglycan layers (34, 35) (Figure 4). This result suggests the cell, during φX174 infection, was experiencing membrane integrity problems, possibly of the peptidoglycan layer, resulting in the up-regulation of these membrane anchoring proteins.

Previous work has demonstrated that phage lysis protein E acts to directly disrupt the activity of integral inner membrane protein MraY (36–38). MraY catalyzes the first membrane associated step of peptidoglycan biosynthesis (39), and subsequent interruption of MraY function by protein E is believed to result in the loss of membrane integrity through synthesis inhibition (37). Interestingly, we saw no up-regulation of the cytoplasmic component of the peptidoglycan biosynthetic pathway that lies immediately upstream of the MraY protein, but did see up-regulation of MraY (FC +1.8) and all observed downstream inner membrane proteins in this pathway: MurG (FC +1.7), and FtsW (FC +1.7) (Figure 5).

**Figure 5:**
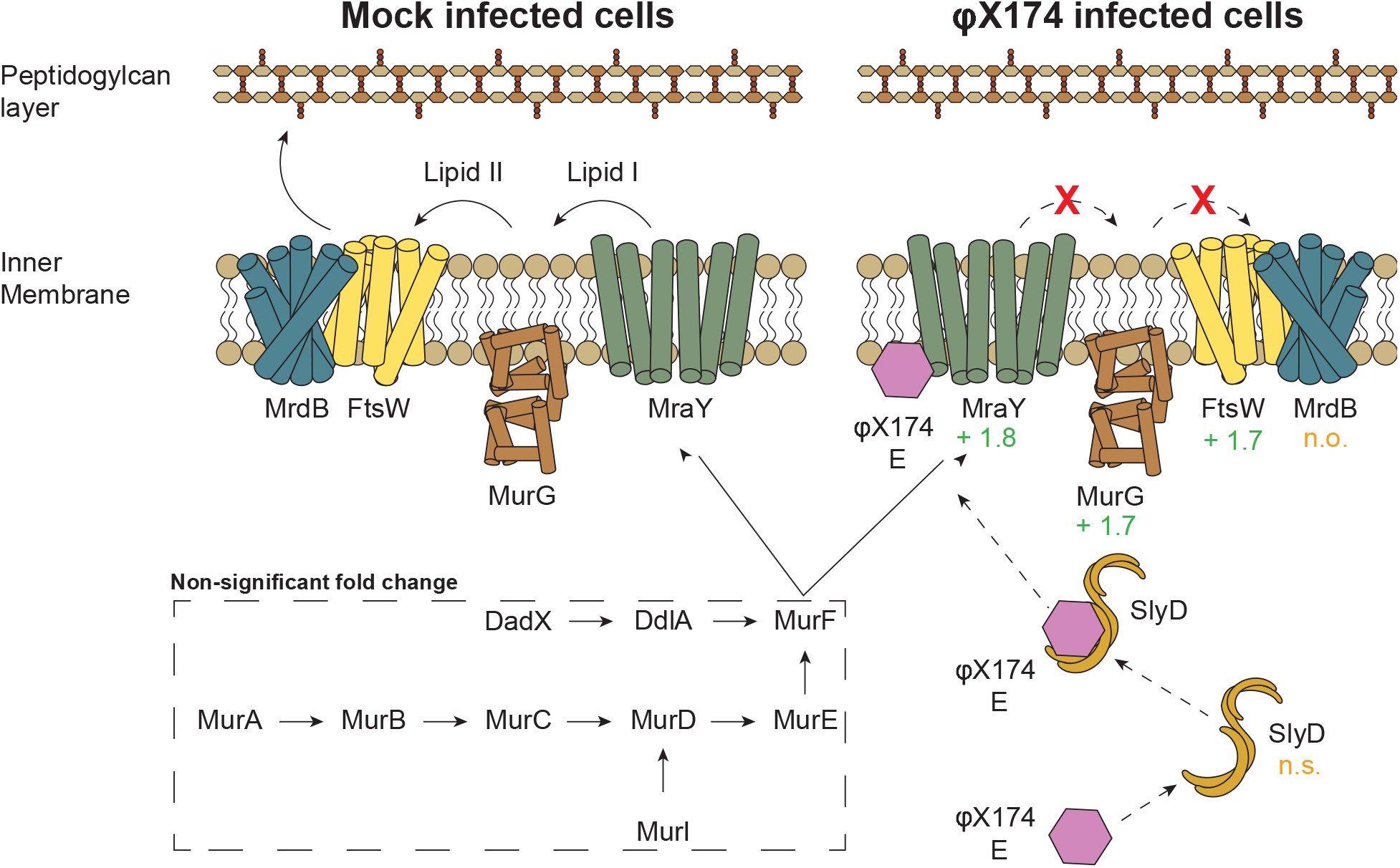
Selective up-regulation of the membrane bound steps of peptidoglycan synthesis observed during infection. The upregulated MraY, MurG, and FtsW are putatively a result of protein E inhibited MraY and subsequent peptidoglycan destabilization due to reduced synthesis. Values indicate protein fold-change during infection (75P/75C protein samples); n.o: protein not observed; n.s: protein observed but not significantly altered in abundance between conditions.

In contrast to the extensive proteome-level response, only 16 differentially expressed RNA transcripts were attributed to COG category M. Within this grouping, only two genes, *murI* and *yjdB*, are involved in lipopolysaccharide biosynthesis. *murI*, which is responsible for lipopolysaccharide core region biosynthesis by catalyzing the racemization of L-glutamate to D-glutamate (40) was down-regulated (FC −1.5) while *eptA* which catalyzes the addition of phosphoethanolamine to the lipid A glucosamine disaccharide (41) was up-regulated (FC +1.6). These results highlight divergence between the captured proteomic and transcriptomic responses.

### Transcriptomics identifies significant up-regulation of the phage-shock protein stress response operon revealing additional cellular response to membrane damage

A major transcriptional response of *E. coli* C122 to φX174 infection was the upregulation of gene products responsible for membrane repair functions and the maintenance of the proton motive force. This was primarily seen through the upregulation of multiple components of the phage shock protein (PSP) response whereby *pspB* (FC +1.7), *pspC* (FC +1.7), and *pspD* (FC +1.8) in the *pspABCDE* operon, and *pspG* (FC +1.8), were up-regulated. Collectively, these genes encode protein products responsible for the binding of PspA to damaged membranes for repair functions. At the proteome-level, significant up-regulation was only observed for PspB (FC +2.1), while PspE was down-regulated, and no change was observed for PspA, PspC, and PspF. Two other genes, *ycjF* and *ycjX*, associated with the PSP response in some species (42), were also upregulated at both the transcriptional and protein levels (RNA FCs +2.4 and +3.0 (Figure S2B); Protein FCs +2.6 and +1.7).

One of the most up-regulated genes during infection was *fxsA* (FC +4.4), which was not identified in the proteomic analysis. Its only ascribed function is to competitively sequester PifA in the cytoplasmic membrane, preventing PifA from performing its host suicide mechanism, known as F exclusion. F exclusion disrupts the bacteriophage T7 lifecycle by arresting all viral and cellular process upon PifA interaction with T7 products gp1.2 and gp10A (43, 44). No annotated version of *pifA* is in the available C122 genomes and it is normally found on the F plasmid, which was not present in the cells in this work.

### φX174 infection significantly alters E. coli metabolic gene expression

To understand the effects of the phage infection on the 58 proteins detected in both Growth and Phage sets (Figure 6D) we classified them based on their three distinct patterns of differential regulation (Figure 6A-C). The first set showed φX174 infection dampening protein expression across 17 genes (Figure 6A). PANTHER GO biological process analysis of this list found enrichment in proteins involved in the tricarboxylic acid cycle (4 proteins, p-value = 5.79E-06), indicative of a disruption to overall metabolic activity during phage infection. The second group of genes showed increased expression in the Phage protein set (Figure 6B). PANTHER GO molecular function analysis on these 14 proteins found enrichment for transporter activity (9 proteins, p-value = 3.59E-06), or more specifically, carboxylic acid transmembrane transporter activity (5 proteins, p-value = 8.50E-06). This is reflective of COG distributions of the entire 58 proteins (Figure 6E), where we see the majority of the highlighted categories being trafficking or transporter groups. The third case showed inverse protein expression pattern between Growth and Phage sets (Figure 6C), which is especially intriguing as it would indicate these proteins are particularly important to phage infection. A PANTHER GO biological process or molecular function analysis of these 28 genes revealed no significant enrichment. However, 15 of the 28 proteins are located in the cell envelope (p-value = 2.82E-11) indicating significant and likely diverse transporter functions from this group of proteins. Of particular interest are aforementioned proteins Lpp and OmpA, as well as OmpA structural homolog, OmpX (FC +3.7) (Figure 4) (45), all of which were found to have large fold-change inductions during phage infection, but were down-regulated during mock-infection (Figure 6C).

**Figure 6:**
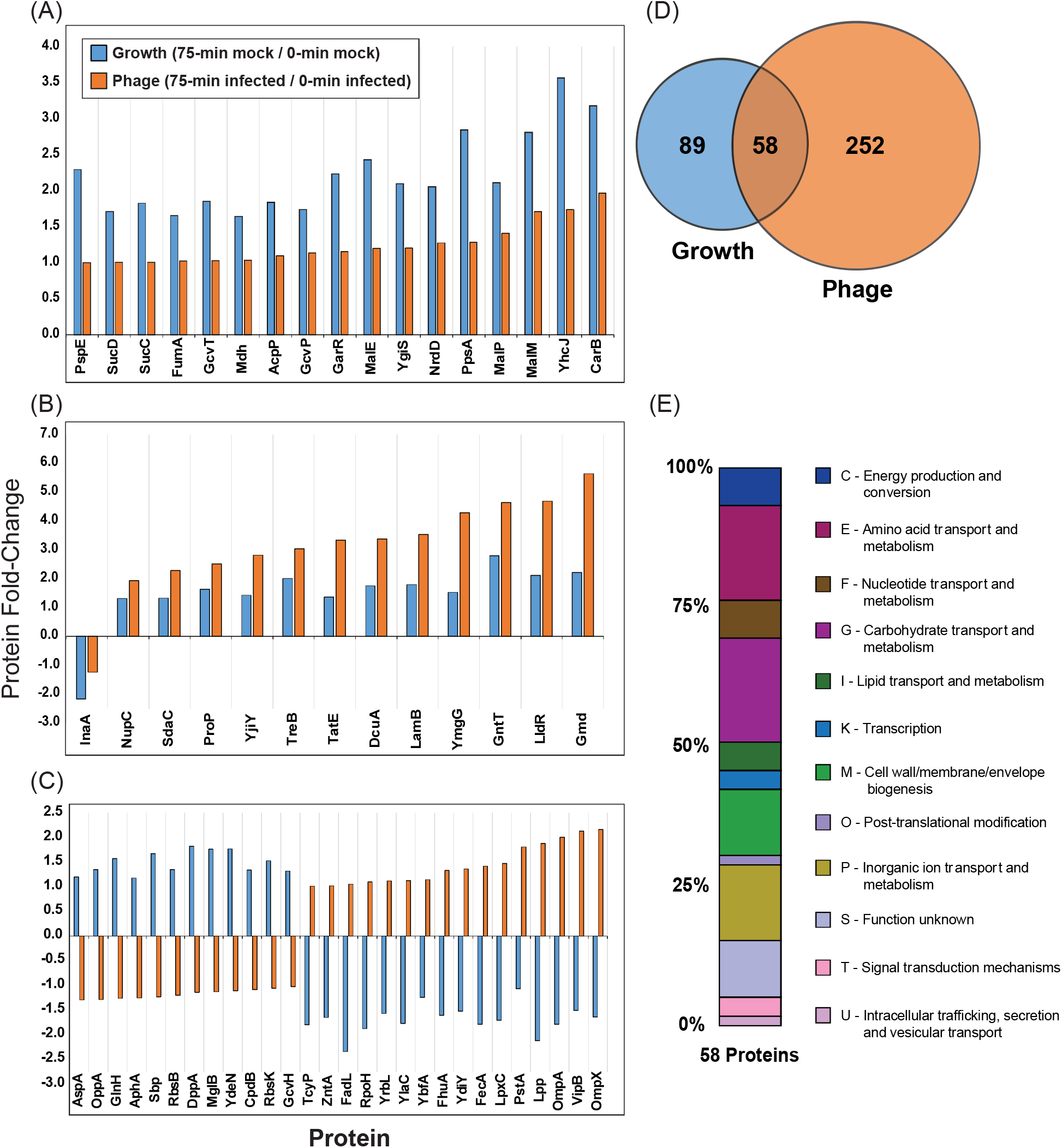
Differential expression patterns of the 58 *E. coli* C122 proteins shared by Growth and Phage sets. Subset of the 58 host proteins where φX174 infection: (A) reduces, (B) increases, (C) inverts expression. (D) Euler plot of all significantly differentially expressed proteins within the Growth and Phage sets. (E) Clusters of orthologous groups (COG) functional categories used to represent major biological functions and their percentage (%) distribution across different groupings for the 58 shared proteins. COGs were assigned to C122 genes using the EggNOG-mapper (23).

Interestingly, OmpA was recently identified as a competitive binder of the lipoprotein RcsF (45), which we also identified as significantly up-regulated during phage infection (FC +1.6). RcsF is a positive regulator of the Rcs stress response (46), of which we identified the significant up-regulation of Rcs components RcsD (FC +1.7) during infection.

Biological process GO term enrichments of the transcriptomics data showed broad representation of two different host functions which affected host metabolic activity when mapped to COG categories. The first of these host functions corresponded to the COG categories nucleoside transport and metabolism (F), amino acid transport and metabolism (E), and carbohydrate transport and metabolism (G). The COG categories were represented with the enriched biological functions of plasma membrane glucose import (17.9 fold, 4 genes, p-value = 4.50E-04), glycine catabolic process (14.3 fold, 4 genes, p-value = 7.77E-04), nucleoside transmembrane transport (11.2 fold, 5 genes, p-value = 3.59E-04), and nucleobase-containing small molecule interconversion (5.5 fold, 7 genes, p-value = 7.54E-04). The second host function mapped to COG categories corresponded to energy production and conversion (C) with the enriched biological functions ATP synthesis coupled proton transport (11.2 fold, 5 genes, p-value = 3.59E-04), gluconeogenesis (7.7 fold, 6 genes, p-value = 4.24E-04), and glycolytic process (6.0 fold, 7 genes, p-value = 4.81E-04) (Figure S4).

### φX174 infection strongly activates heat shock chaperone response

The strongest host response we observed at both the protein and RNA level was the heat shock response, led by the small heat shock proteins (sHSPs) IbpA and IbpB (Figure 1). Our proteomics and transcriptomic data showed that they had the largest increase in expression of any host gene products (Figure 1C). Both *ibpA* and *ibpB* are controlled by the σ^32^ regulon (47) and their large up-regulation during φX174 infection is consistent with the significant increase in RpoH transcription factor σ^32^ (FC +2.0), along with chaperones ClpB (FC +1.7) and DnaJ (FC +1.9) (Figure 7). Notably, only a subset of the σ^32^ heat-shock stress response regulon was activated during φX174 infection, as we observed only 11 proteins and 17 transcripts belonging to this regulon differentially regulated (Figure 7, and Supplementary File S3). Similarly, the general stress response master regulator, σ^38^ (RpoS) was not differentially regulated and only a small proportion of proteins (5%) from the genes under its control were differentially regulated during phage infection (Supplementary File S3).

**Figure 7:**
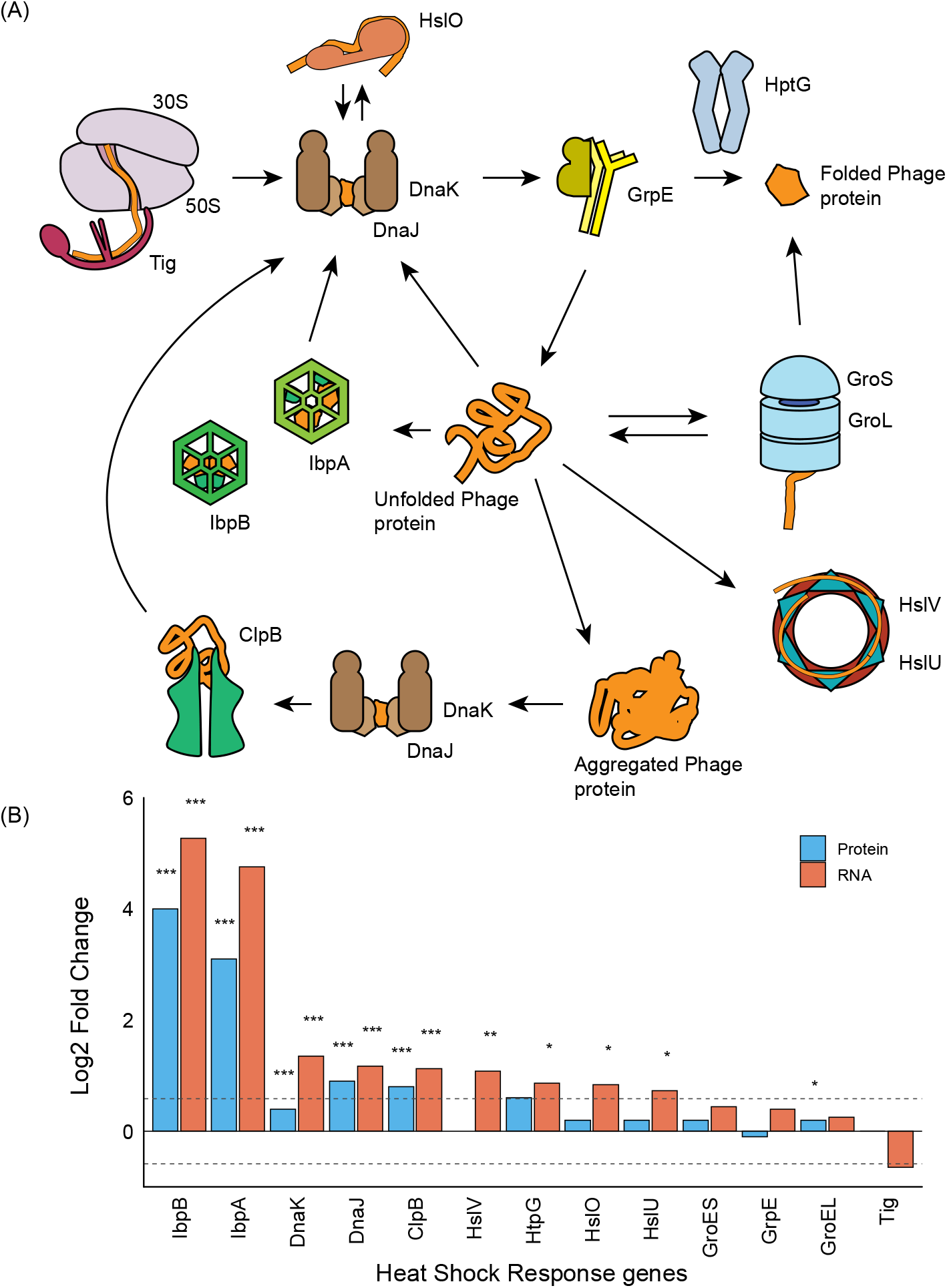
Large changes to gene expression across the heat shock chaperone and protein folding pathway in *E. coli* C122 during φX174 infection. Proteins depicted in this pathway are transcriptionally regulated by transcription factor RpoH (σ^32^), which we saw up-regulated during infection. (A) The chaperone system revealing the process controlling protein folding from synthesis to native protein state. The chaperone trigger factor (Tig) binds to the translating ribosome and the growing peptide chain. The DnaK/J chaperone proteins aid in folding by binding to hydrophobic regions of nascent proteins preventing their misfolding and aggregation. DnaK/J may bind to HtpG to reactivate inactive protein substrates on the path to native protein fold (48). The holdase HslO binds to unfolded protein which is released to DnaK for folding (49). GrpE may interact with DnaK thereby dissociating ADP and releasing the bound protein (50). Small heat shock chaperones IbpA/B bind nascent proteins preventing their irreversible aggregation into inclusion bodies, and await the availability of folding chaperones DnaK/J and GroEL/ES. The main folding machine of the system is the GroEL/ES complex. This ATP-dependent system facilitates folding of nascent proteins within the complex’s inner surface releasing proteins in their native state. In the case of protein aggregates produced from improperly or partially folded proteins, the chaperone ClpB mediates their disaggregation for their re-folding. Unfolded protein may also shuffle to the HslUV protease/ATPase complex for proteolysis (51). (B) Differential expression of proteins and transcripts measured at lysis. Dashed lines indicate the Log2 fold change ± 0.58 values used as significance criteria. P-values are shown as *** for ≤ 0.001, ** for ≤ 0.01, and * as ≤ 0.05.

Together, these data indicate that φX174 infection was not inducing a general stress response via the σ^38^ or σ^32^ regulon, and that there was selective up-regulation of members of the heat shock chaperone network (Figure 7).

## DISCUSSION

In this work, we have comprehensively measured the *E. coli* C122 host response to φX174 infection at the proteomic and transcriptomic level, representing the first characterization of the dynamics of *Microviridae* infection at this level of detail. We identified 2,184 proteins and quantified all 11 phage proteins and 1,752 host proteins across seven time-points spanning the entire infection cycle. Transcriptomics identified significant changes during infection including differential expression of 528 genes, constituting 11 % of the total transcriptome, similar to other studies (52).

We found no evidence for substantial host proteome remodeling during the early stages of infection prior to 60-minutes (Figure 3A), which aligns with previous work showing φX174 has a minimal impact on host protein production until just prior to lysis (53). It is well-established that phage infection can cause drastic and broad metabolic remodeling of infected cells (11, 54, 55), which is in agreement with what we saw for both our proteomic and transcriptomic datasets.

We showed that membrane damage responses dominated the host response to φX174 infection (Figures 4 and 5). We also found membrane lipoprotein maturation and transportation pathway enrichment (Figure 4) which suggests an up-regulation of this pathway in response to the host’s need to traffic certain lipoproteins to the outer membrane, such as major lipoprotein (Lpp). Lpp is one of the most abundant (lipo)proteins of *E. coli* (56) and is reliant on the Lol pathway for outer membrane insertion (33). We saw Lpp down-regulated during growth (FC −2.1), and up-regulated during φX174 infection (FC +1.9), thus emphasizing an infection mediated response. Lpp has recently been shown to regulate cell shape and mechanical rigidity by tethering the outer membrane to the peptidoglycan layer (32), which leads us to suggest that during φX174 infection, Lpp is up-regulated to maintain the cell’s membrane integrity.

The most likely model that fits our observations would have lysis protein E from φX174 binding to and inhibiting MraY (36, 37) and causing a decrease in peptidoglycan synthesis which is sensed by the cell and results in upregulation of MraY and inner membrane pathway partners, MurG and FtsW (Figure 5). The cytoplasmic components of the peptidoglycan pathway and chaperone protein, SlyD, which stabilizes the φX174 lysis protein E and is thought to promote its interaction with the inner membrane (57) are not upregulated (Figure 5).

Secondarily, as the φX174-infected *E. coli* cell continues to grow, but is unable to synthesize enough peptidoglycan, the stability of the envelope becomes compromised. The cell senses this and attempts to compensate by upregulating the Lpp, OmpA, and Pal proteins (Figure 4). OmpA also seems to have a role in maintaining envelope integrity by linking the outer-membrane to the peptidoglycan layer through the non-covalent linkage of its C-terminal domain to peptidoglycan (35). In addition to this activity, OmpA has a direct interaction with peptidoglycan stress sensory protein RcsF (45), which we also saw up-regulated during infection (FC +1.6). Pal protein is also a mediator of outer membrane and peptidoglycan stability by forming a structural linkage between both layers (34).

Host membrane responses to phage infection are common and can involve responses such as induction of polysaccharide capsule biosynthesis (58) and phage shock response (55). We saw the significant upregulation of components *pspB*, pspC, *pspG*, and the potentially related genes *ycjX* and *ycjF* (Figure S2), presumably in response to the loss of envelope structural integrity during the lead up to lysis. A similar induction of the phage shock response was observed with phage PRD1 infection of *E. coli* (59) and antibiotics that inhibit peptidoglycan synthesis (60).

Both proteomics and transcriptomics showed a massive up-regulation of small heat shock proteins IbpA (FC +8.7) and IbpB (FC +15.6) and their transcripts (FC +27.1 and +38.6, respectively) during infection. The IbpA/B proteins are well-described and the sHSP family is conserved across all domains of life (61). They are typically described as “holdase” proteins, as they function as chaperones binding mis-folded or un-folded (nascent) proteins to prevent their irreversible aggregation within the cytosol, while they wait for available ATP-mediated folding machinery (62, 63) (Figure 7). The large mismatch between *ibpA/B* RNA and protein fold-changes in our measurements may be due to the RNA thermometer function encoded within their transcripts that ensures they are only maximally translated at elevated temperatures (64).

We speculate that the up-regulation of IbpA/B proteins is a conserved nonspecific bacterial response to viral infection based upon their up-regulation during both phage infection (59, 65) as well as during heterologous protein overexpression (66, 67). Therefore, as a result of their currently defined function (62, 63), IbpA/B up-regulation in *E. coli* C122 during φX174 infection is a host response to higher than normal levels of protein production (in this case, phage protein production), and in the absence of available protein folding machinery, irreversible protein aggregation or mis-folding is prevented through the up-regulation of these holdase proteins.

Phage may also benefit from the specific features of the IbpA/B function by having a queue of proteins held in a near-folded state that can be quickly folded into their final forms by HSP70 (DnaK) and HSP100 (ClpB) activity. Proteins bound by IbpA/B are not spontaneously released and can only be released by DnaK and ClpB (64), introducing a buffering system and control point for virion assembly.

The *ibpA/B* genes have never been identified from genetic screens of host-factors necessary for capsid morphogenesis, but sHSPs are encoded in the genomes of cyanophage infecting *Synechococcus* and *Prochlorococcus* groups (68, 69). Furthermore, a bacteriophage-encoded J-domain protein (Rki) has been identified in T4-related enterobacteriophage RB43 that interacts with *E. coli* J-domain interacting chaperone protein DnaK (HSP70) and stabilizes the σ^32^ heat-shock response (70). MS2 phage relies on DnaJ (HSP40) (71), while the HSP60 chaperones GroEL/ES are required for phages λ, PRD1, HK97, and T7 morphogenesis (72–75). GroEL-like proteins have also been found encoded within newly annotated bacteriophage genomes (76). In *groEL/groES* mutants phage T4 head morphogenesis is disrupted and results in a random aggregation of phage head proteins attached to the inner membranes (77), pointing to the critical disaggregating and/or folding role of the HSP-chaperones in phage capsid assembly. Young, et al (1989) found that lysis sensitivity increased in *E. coli* when plasmid encoded φX174 lysis protein E and heat-shock genes *dnaK, dnaJ, groEL*, and *grpE* are present (78). Recently, φX174 has been suggested to use heat shock promoters to co-opt host responses to infection to drive its lifecycle (12, 79). Therefore, together with our data, we propose that the host heat-shock response, and in particular IbpA/B, are important components for φX174 replication.

## MATERIALS AND METHODS

### φX174 infection

Methods are composed in accordance with (80). Media and buffer components from Sigma Aldrich, unless stated otherwise. *E. coli* C122 (Public Health England NCTC122) was grown overnight at 37°C/250 RPM in phage-LB (81), then inoculated 1/100 in fresh phage-LB and grown to mid-log phase before addition of phage (37 °C/250 RPM). Wild-type φX174 (14) was used for all infections of *E. coli* C122 at an MOI = 5. Phage infection synchronized by pelleting mid-log cultures (4,000 RCF/8-min) and resuspending the pellet to 1/10^th^ of the original growth volume with cold HFB buffer (82). Starved cells were incubated 16 °C/30-min to facilitate phage attachment and to prevent DNA ejection (83). After 30-min, infection was initiated with the addition of phage-LB (37 °C) to the original volume. Controls (mock-infected) had the same volume of phage-LB and 10 % (v/v) glycerol added at mid-log as the phage-infected samples.

### Protein purification and tandem mass tagging (TMT) labelling

*E. coli* C122 was grown to mid-log in 200 mL culture volume (37 °C/250 RPM) then split into 2×100 mL volumes: one to be infected with φX174 at MOI = 5, and the other mock-infected with equal volume phage-LB with 10 % (v/v) glycerol (storage solution of φX174). Culture growth was synchronized with 10 mL of ice-cold HFB, and infection was initiated with 90 mL phage-LB (37 °C). At 0, 15, 30, 60, and 75-min postinfection, 12 mL was removed from each culture (without replacement) and placed on ice. At the conclusion of the time-course, intact cells were pelleted 3,220 RCF/8-min. The samples were then suspended in cold 1X phosphate-buffered saline (PBS) and washed an additional two times, discarding the supernatant. Cell pellets were stored overnight at −20 °C. Cell pellets were suspended in 500 μL lysis buffer (100 mM Tris-HCl, 1 % (v/v) sodium dodecyl sulfate, 8 M urea, 1X protease inhibitor cocktail solution (Roche, Switzerland). Reconstituted cells were lysed with an ultrasonic probe with 15-pulses at 30 % amplitude. Samples were then pelleted 6,000 RCF/10-min and supernatant moved to new tube. Proteins were reduced 37 °C/1-hr in 10 mM dithiothreitol, followed by alkylation with 30 mM iodoacetamide for 1-hr in dark. Residual iodoacetamide quenched with an equal molarity of dithiothreitol.

Proteins were precipitated by addition of 2 mL methanol, 0.5 mL of chloroform, and 2 mL of water (all ice cold). Samples were pelleted 5,000 RCF/10-min and protein pellet removed, washed with additional methanol, and dried. Protein pellets were solubilized with 100 mM Tris-HCl/8 M urea. Solutions were pelleted 12,000 RCF/10-min and supernatant collected in new tube. Samples were diluted to urea concentration of 1.6 M through addition of 100 mM Tris-HCl, followed by protein quantitation (Pierce™ BCA Protein Assay Kit, ThermoFisher Scientific). 150 μg of protein from each sample digested overnight/37 °C with 1.5 μg trypsin (Promega), followed by addition of further 1.5 μg of trypsin and digestion for an additional 4-hrs. Peptides were acidified with formic acid at 1 % (v/v) and then C18 stage-tip purified (84). Peptide samples were dried down using vacuum centrifugation, followed by resuspension in 200 mM HEPES buffer (pH 8.8, adjusted with NaOH). Peptides were quantified (Micro BCA™ Protein Assay Kit, ThermoFisher Scientific), and 35 μg peptide from each sample brought up to 140 μL using HEPES buffer. Normalization controls were produced through pooling mock-infected samples (3 μg of peptide per mock-infected sample) – designated N.C, and infected samples (3 μg of peptide per infected sample) – designated N.P.

Chemical labelling of peptide samples was performed using TMT10Plex™ Isobaric Label Reagent Set (ThermoFisher Scientific, USA). 0.2 mg of TMT tag was used to label each sample (Table S1). Note: 128C tag was reserved for N. C. 11.25 μg of N.C was labelled per channel. Similarly, 131 tag was reserved for N.P. 5.375 μg of N.P was labelled per channel. After labelling, 8 μL of 5 % (v/v) hydroxylamine was added to each sample to quench residual tags. Lastly, samples were combined across channels, dried down using vacuum centrifugation, and then purified as described (84).

### High pH peptide pre-fractionation

Dried peptide samples (TMT channels 1-4) were suspended in high-pH buffer (5 mM NH_4_OH, pH 10.5), and loaded onto Agilent ZORBAX Extend-C18 column (3.5 μM bead size, 300 Å pore size, 2.1 mM x 150 mM), washed with buffer A (0.1 % (v/v) formic acid, 2 % (v/v) acetonitrile) for 10-min, followed by elution with increasing gradient of buffer B (5 mM NH_4_OH, 90 % (v/v) acetonitrile). The gradient of buffer B was 3 % to 30 % for 55-min then to 70 % for 10-min, and finally to 90 % for 5-min at a flow-rate of 300 μL/minute. Samples were collected every minute and pooled into 13 fractions, vacuum centrifuged, and then suspended in buffer A to concentration of 0.1 μg/μL.

### Mass spectrometry

Samples were analyzed on a Q-Exactive mass spectrometer (ThermoFisher Scientific) coupled to EASY-nLC1000 system (ThermoFisher Scientific, USA). Peptide samples were injected onto the LC system using buffer A and were bound on a 75 μM x 100 mM C18 HALO column (2.7 μM bead size, 160 Å pore size). A flow rate of 300 nL/minute using an increasing linear gradient of buffer B (0.1 % (v/v) formic acid, 99.9 % (v/v) acetonitrile) was run from 1 % to 30 % for 110-min followed by 85 % buffer B for 10-min. The mass spectrometer was operated in top-10 mode, with a full scan set at a resolution of 70,000 (at 400 *m/z*) across the *m/z* range of 350-1850 (isolation window of 0.7 *m/z*), and an automatic gains control (AGC) target of 1e6 (or maximum fill time of 250 ms). Selected pre-cursor ions were transferred from the C-trap to the higher energy collision dissociation (HCD) cell for fragmentation at a normalized collision energy of 35 % with precursor dynamic exclusion of 90-sec. MS/MS spectra was collected at a resolution of 70,000 (at 200 *m/z*) with an AGC of 1e5, maximum injection time of 250 ms and a fixed first mass of 115.0 *m/z*.

Raw files exported to Proteome Discoverer v2.1 (ThermoFisher Scientific) for processing. Precursors selected for fragmentation that had greater than 30 % interference were excluded from analysis. Methionine oxidation, N-terminal carbamylation, asparagine and glutamine deamidation, N-terminal acetylation, N-terminal glutamic acid to N-pyro-glutamine, N-terminal glutamine to N-pyro-glutamine, and TMT10plex labelling of primary amines were selected as dynamic modifications. Cysteine carbamidomethylation was designated as a fixed modification. Minimum peptide length was set at 5. Spectra were searched against a custom *E. coli* C122 database that included the phage proteins, totaling 3,806 entries (Supplemental File 4). False discovery rates were fixed 1 % at the peptide and protein level. Runs were normalized by dividing by all channels (e.g. 126, 128N) by the pooled control (128C), log_2_ transformed and further normalized by subtracting by the median of each sample group (Table S1). Sample infected 60_1_ was excluded from statistical tests due to phage proteins being designated as outliers. Student’s t-test was performed on the log_2_ transformed data. Due to effects of ratio compression and to appropriately control for false-discovery in a (low-power) medium-scale experiment (85), significance was deemed with protein fold-change cut-of ≥ ±1.5, p-value <0.025.

### RNA isolation and RNA-seq

*E. coli* C122 was grown and infected with wild-type φX174 at MOI = 5, 5 mL samples removed from infected and mock replicates at initial infection and at lysis, pelleted (3,500 RCF/5-min/4 °C), and resuspended in 200 μL 1X PBS. 400 μL of RNAprotect bacterial reagent (Qiagen: 76506) added, vortexed, and incubated 10-min/room temperature. Following protection, RNeasy Mini Kit (Qiagen: 74106) used with optional DNAse step. RNA samples were then eluted, RNA quantified (Qubit RNA HS Assay Kit, ThermoFisher Scientific: Q32852), and stored at −80 °C. Sequencing library preparation performed by Macrogen Inc. (South Korea). The rRNA from samples were depleted with a Ribo-Zero Kit (Illumina) and the RNA library generated with a TruSeq Stranded mRNA kit for microbes. Libraries sequenced with Illumina HiSeq 2500 (2×100bp mode). PhiX Control Library not used during the sequencing run to avoid cross-contamination.

Read quality checked with FastQC followed by Rsubread analysis using standard protocol (86). BAM files were processed with Rsubread featureCount function against a manually generated General Feature Format (GTF) file derived from the NZ_LT906474.1 RefSeq GFF3 file. RNA differential expression was analyzed using DESeq2 (87) and visualized using ggplot2 R package (88). To analyze trends in differential expression, gene names were converted to UniProt identifiers using the Retrieve/ID mapping tool (89) and searched against the PANTHER database Gene list analysis tool (90) using the statistical over-representation test against the internal reference *Escherichia coli* gene list to generate overrepresented GO Terms for both cellular compartments and biological processes. The converted UniProt IDs were used to extract individual GO Terms as before for all genes in the differentially expressed bins using the EcoCyc database (91).

### Databases and gene analysis

*E. coli* C122 gene annotations for proteomics experiment and COG assignments were produced through the EggNOG mapper using default parameters (23). For σ^32^ and σ^38^ transcription factor assignments, we extracted orthologous gene members from the gold-standard *E. coli* K12 gene annotations from the EcoCyc database (91), and assigned matching annotations to C122 genes (Supplemental File S3). The PANTHER (24) database’s statistical over-representation tests and functional classifications was used to map protein and transcript enrichments.

Mass spectrometry proteomics data have been deposited to the ProteomeXchange Consortium via the PRIDE (92) partner repository with the dataset identifier PXD021681.

## SUPPLEMENTAL MATERIAL

Supplementary File S1: TMT proteomics data

Supplementary file S2: Transcriptional dataset of all C122 genes

Supplementary File S3: Sigma32 and Sigma38 analysis

Supplementary File S4: MS protein database

## ACKNOWLEDGEMENTS

We recognize that this research was conducted on the traditional lands of the Wattamattagal clan of the Darug nation. BWW was supported by a Macquarie Research Excellence PhD Scholarship. We thank Dr. Karthik Kamath, Dr. David Cantor, Sasha Tetu, and Varsha Naidu for assistance. Aspects of this research were conducted at the Australian Proteome Analysis Facility. PRJ was supported by the Molecular Sciences Department, Faculty of Science & Engineering, and the Deputy Vice-Chancellor (Research) of Macquarie University. DYL is a recipient of the Macquarie University Research Excellence PhD scholarship (MQRES) and CSIRO PhD Scholarship Program in Synthetic Biology (Synthetic Biology Future Science Platform). BWW was supported by a Macquarie Research Excellence PhD Scholarship.

The contributions of author of this work according to the CRediT contribution taxonomy was: BWW: Conceptualization, Formal analysis, Investigation, Methodology, Visualization, Writing – Original draft, Writing – review & editing; DYL: Formal analysis, Investigation, Methodology, Visualization, Writing - original draft, Writing - review & editing; MM: Investigation, Methodology; DP: Formal analysis, Methodology; MPM: Funding acquisition; Project administration; Resources; Supervision; Writing - review & editing. PRJ: Conceptualization; Formal analysis; Funding acquisition; Methodology; Project administration; Resources; Supervision; Writing - review & editing.

## Notes

### Competing Interest Statement

The authors have declared no competing interest.

