## Supplementary for "Proteomic and transcriptomic analysis of *Microviridae* φXI74 infection reveals broad up-regulation of host membrane damage and heat shock responses"

**CONTENTS**

FIGURES

Figure S1: Mock and φX174 infected *E. coli* C122 Analysis.

Figure S2: Differential expression of φX174 infected NCTC122 genes.

Figure S3: Visualization of the major biological functions of the differentially expressed *E. coli* C122 genes through their COG distributions.

Figure S4: RNA-seq differentially expressed E. coli C122 gene major cellular localization GO terms

Figure S5: RNA-seq differentially expressed E. coli C122 gene major biological function GO terms

TABLE

Table S1: TMT Labelling scheme

FILES

Supplementary File S1: TMT proteomics data

Supplementary file S2: Transcriptional dataset of all C122 genes

Supplementary File S3: Sigma32 and Sigma38 analysis

Supplementary File S4: MS protein database

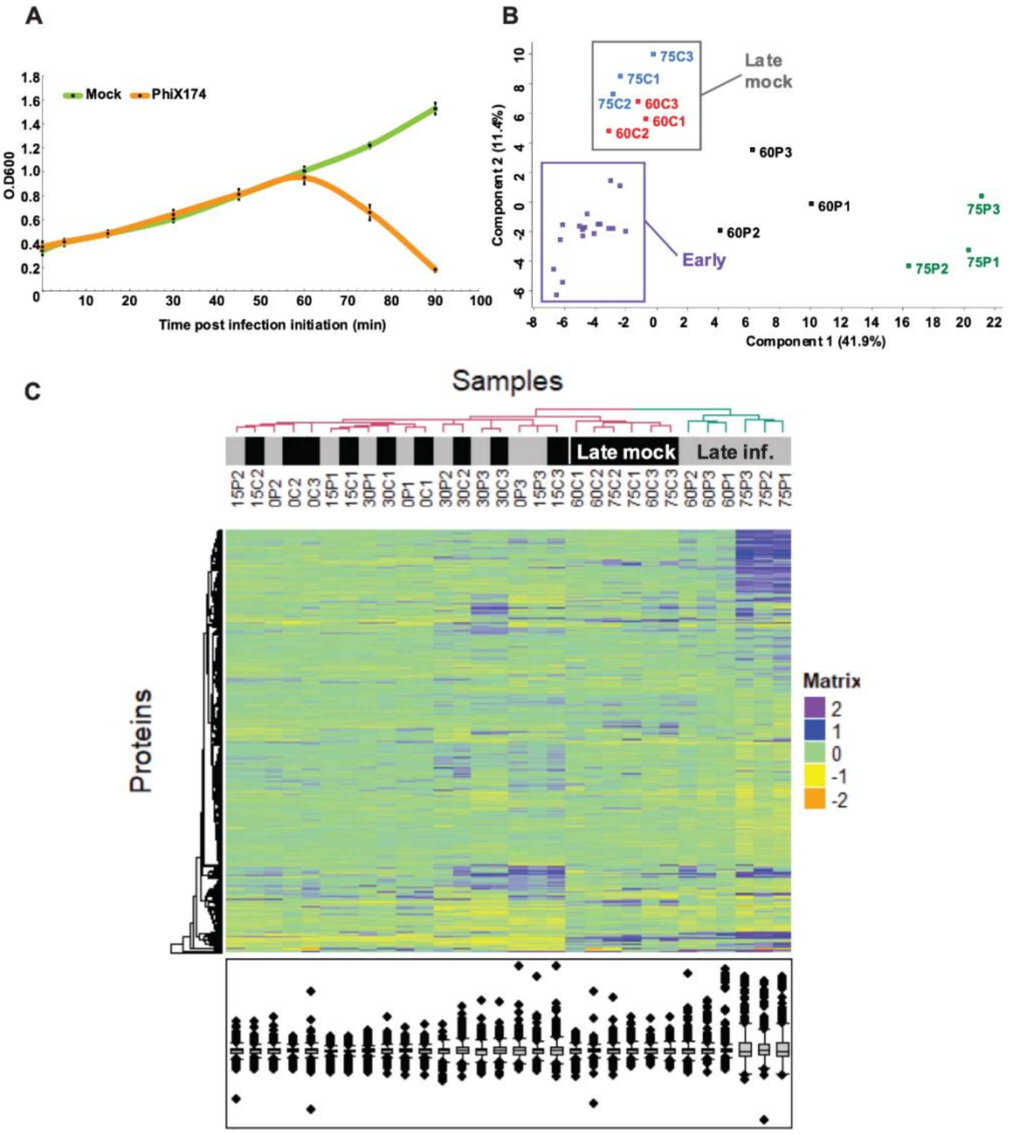


**Figure S1 Mock and φX174 infected *E. coli* C122 Analysis. (A)** Lysis curve of φX174 and *E. coli* C122. Lysis was observed at 60-minutes post-infection initiation. **(B)** 2-Dimensional principle component analysis (PCA) of mock-infected (C) and φX174-infected (P) samples. Separate clustering of the later time-point control samples (Late mock) to that of the early time-points of both mock-infected and φX174-infected (Early) can be observed. Similarly, separate clustering of later time points of the 60-minute and 75-minute φX174-infected samples (60P and 75P) to each other and of the late mock and early groupings is observed. **(C)** Hierarchical clustering with Euclidean distance of quantified proteins. Clustering of later time-points (late), and their associated condition (mock = mock-infected, inf. = φX174-infected) highlights the change in the proteome over time and by φX174 infection.

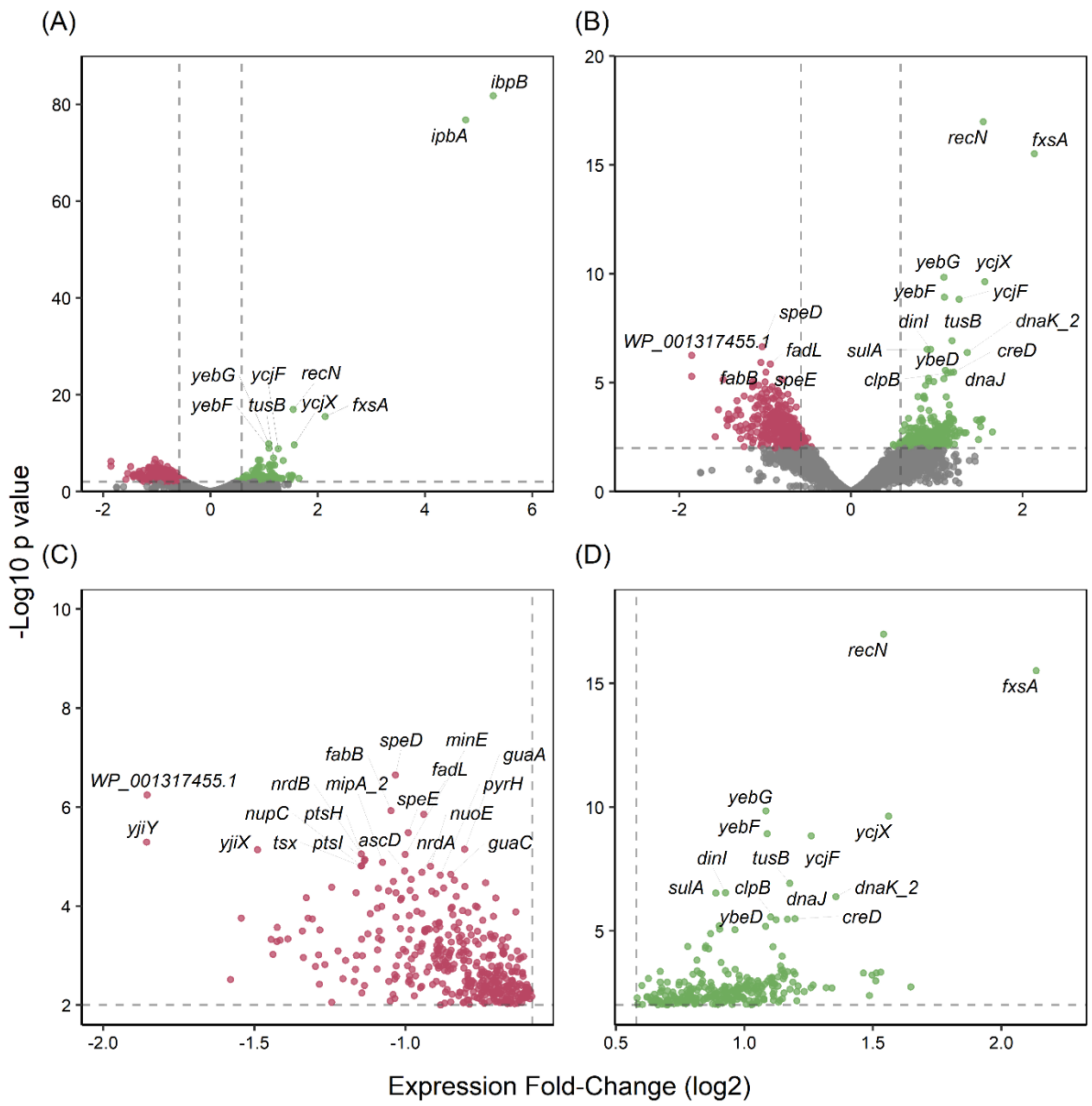


**Figure S2. Differential expression *E. coli* C122 genes during φX174 infection.** **(A)** Differential expression of all genes within infected *E. coli* C122 with significantly up-regulated genes shown in green and significantly down-regulated genes shown in red. Dashed lines represent significance criteria (Log2 fold change = ± 0.585 and p values < 0.05). **(B)** Differentially expressed genes excluding highly up-regulated genes *ibpA* and *ibpB*. **(C)** Down-regulated genes (close up). **(D)** Up-regulated genes (close up). Comparison were made between RNA-sequencing outputs from cultures harvested at lysis in the infected and mock infected samples, samples were analyzed using Rsubread and DESeq2 in biological triplicates.

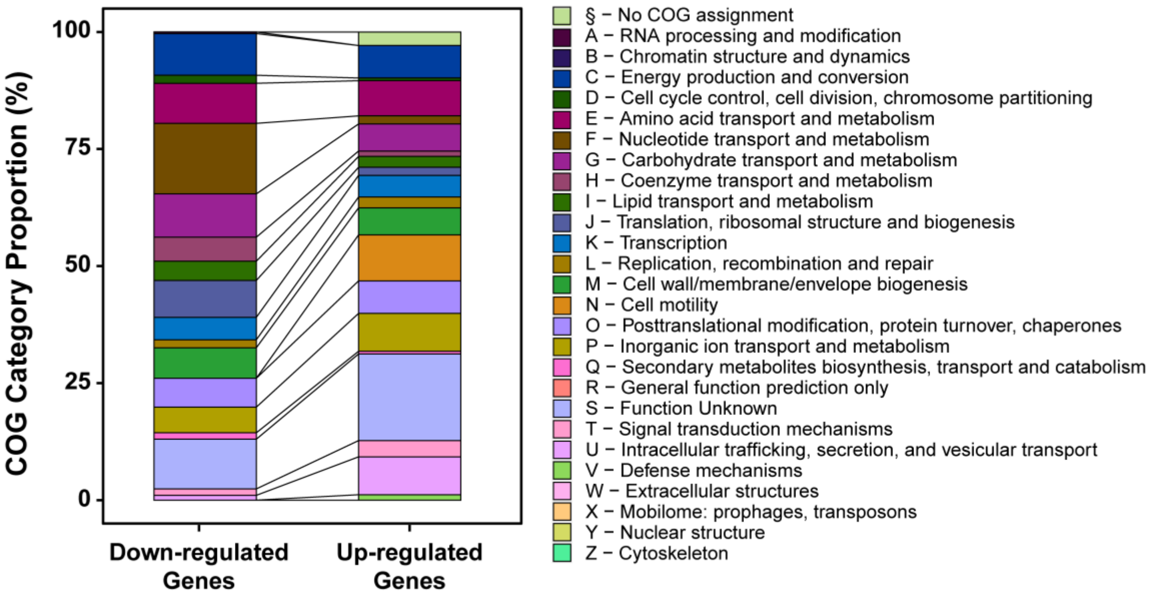


**Figure S3 Biological function COG distributions for differentially expressed *E. coli* C122 genes in the RNA-seq dataset.** The enrichment of COG terms in the down- and up-regulated RNA-seq datasets from C122 at lysis. UniProt IDs mapped to COG terms using EggNOG-Mapper (1). Gene ID with multiple COG terms were converted to multiple single term entries for analysis.

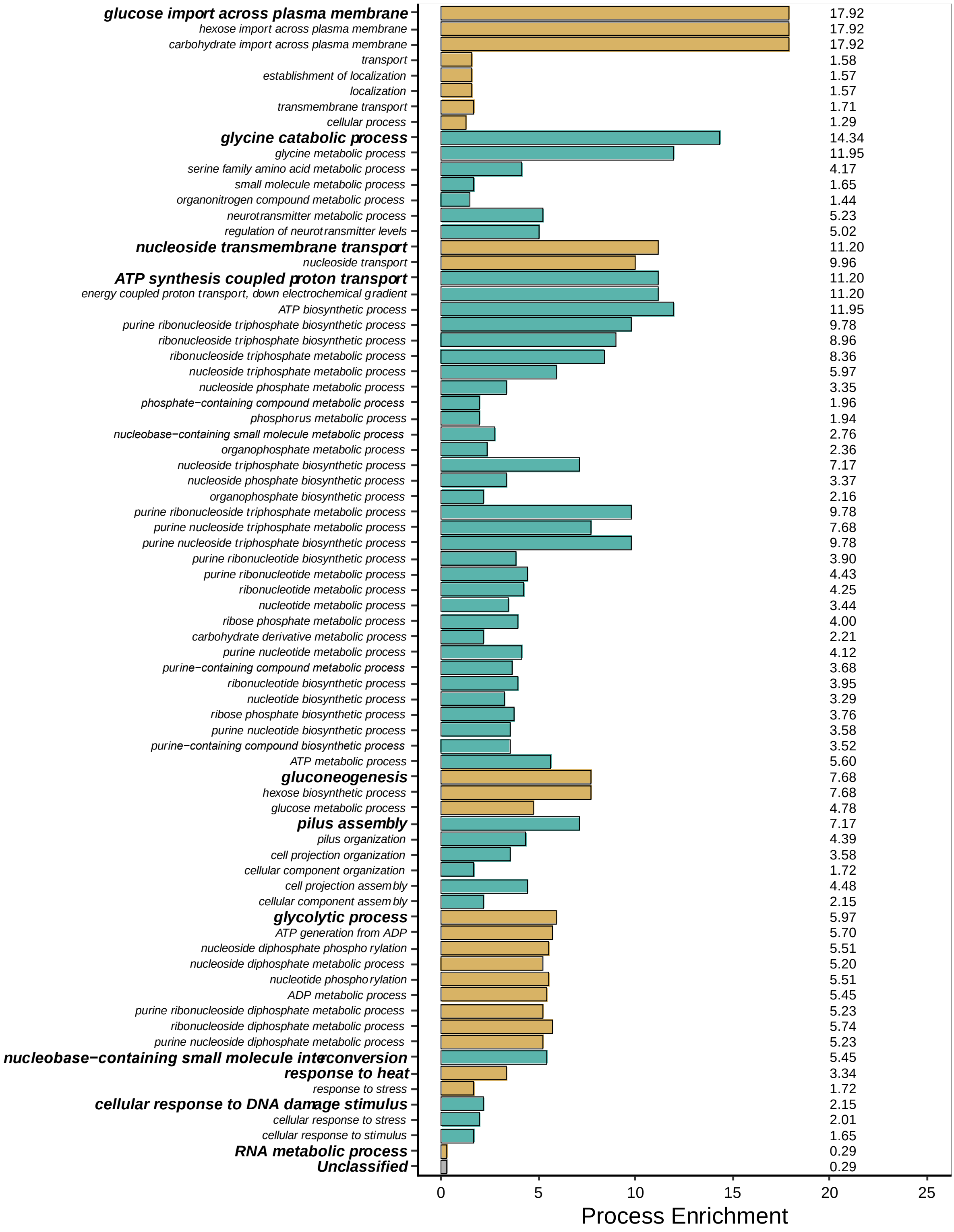


**Figure S4 Major biological function GO terms differentially expressed in RNA-seq dataset.** Differentially expressed genes from *E. coli* C122 were analyzed using the PANTHER over-representation test using the GO terms for Biological Function (2). All presented data displayed a p value of < 0.01.

**Table S1**: **TMT Labelling scheme.** There were four TMT 10-plex experiments with each channel comprising a different sample, except for the 128C and 131 channels. Channel 128C contained the pooled mock-infected samples (designated N.C), and channel 131 contained the pooled φX174-infected samples (designated N.P)

| | Mock-infected | | | | | ϕX174-infected | | | | |
|---|---|---|---|---|---|---|---|---|---|---|
| **TMT** | **126** | **127N** | **127C** | **128N** | **128C** | **129N** | **129C** | **130N** | **130C** | **131** |
| **1** | $0_1$ | $15_1$ | $30_1$ | $60_1$ | N.C | $0_1$ | $15_1$ | $30_1$ | $60_1$ | N.P |
| **2** | $75_1$ | $0_2$ | $15_2$ | $30_2$ | N.C | $75_1$ | $0_2$ | $15_2$ | $30_2$ | N.P |
| **3** | $60_2$ | $75_2$ | $0_3$ | $15_3$ | N.C | $60_2$ | $75_2$ | $0_3$ | $15_3$ | N.P |
| **4** | $30_3$ | $60_3$ | $75_3$ | | N.C | $30_3$ | $60_3$ | $75_3$ | | N.P |

Subscript = replicate number
